# ELAV mediates circular RNA biogenesis in neurons

**DOI:** 10.1101/2024.06.22.600180

**Authors:** Carlos Alfonso-Gonzalez, Sarah Holec, Sakshi Gorey, Mengjin Shi, Michael Rauer, Judit Carrasco, Stylianos Tsagkris, Valérie Hilgers

**Affiliations:** Max Planck Institute of Immunobiology and Epigenetics, Freiburg, Germany; Faculty of Biology, University of Freiburg, Freiburg, Germany; Signalling Research Centre CIBSS, University of Freiburg, Schänzlestraße 18, 79104 Freiburg, Germany; Institut für Medizinische Bioinformatik und Systemmedizin, University of Freiburg, Freiburg, Germany; Discovery Sciences, BioPharmaceuticals R&D, AstraZeneca, CB2 0AA, Cambridge, United Kingdom; Epigenetics & Neurobiology Unit, European Molecular Biology Laboratory (EMBL)-Rome, Adriano Buzzati-Traverso Campus, Rome, Italy

**Keywords:** circular RNAs, nervous system, RNA-binding proteins, ELAV, Drosophila, splicing, back-splicing, reverse complementary matches (RCMs), RNA processing, RNA binding motif

## Abstract

Circular RNAs (circRNAs) arise from back-splicing of precursor RNAs and accumulate in the nervous system of animals, where they are thought to regulate gene expression and synaptic function. Here, we show that neuronal circRNA biosynthesis is mediated by the pan-neuronal RNA-binding protein ELAV. In Drosophila embryos, we characterize the circRNA landscape in normal and *elav* mutant neurons. We find that neuronal circRNAs are globally (>75%) depleted upon ELAV knockout, and induction of ELAV expression drives ectopic RNA circularization. In brain tissue, ELAV binds to pre-mRNA introns flanking putative circRNAs and decreases efficiency of linear splicing in favor of intron pairing at reverse complementary matches, inducing circularization. Together, our data demonstrate that ELAV directly modulates splicing decisions to generate the neuronal circRNA landscape.

## Introduction

Circular RNAs (circRNAs) are single-stranded RNA molecules covalently bound into a continuous loop, found ubiquitously across all domains of life ^1^. In animals, circRNAs are expressed in tissue- and developmental stage-specific patterns in animals, with a particularly high abundance in the nervous system ^2–5^. CircRNAs typically arise from genes linked to synaptic functions. Although their physiological functions remain much less well studied compared to the linear version of the gene, circRNAs have been shown to play important roles in various aspects of brain development and functionality, including neural stemness and neurodegeneration during aging, cognition, and plasticity ^6–11^. For instance, the loss of Cdr1as circRNA disrupts sensorimotor gating, a phenotype associated with several human neuropsychiatric disorders ^12,13^. Moreover, circRNAs expression is altered in multiple neuropathological conditions such as addiction and neurodegeneration, making them potential markers for these diseases ^14,15^. Mechanisms of action differ for distinct circRNAs ^1^ ; they may act as regulators of individual genes ^16^, function globally as sponges for microRNAs and RNA-binding proteins (RBPs) ^17,18^, while others are translated into a functional protein ^19,20^.

CircRNA formation is a co-transcriptional process in which a 5’ splice site is ligated “back” to a 3’ splice site located upstream, thereby forming a circular structure with a characteristic 3’-5’ phosphodiester bond at the back-splicing junction site (BSJ) ^21,22^. Intron pairing is facilitated by co-transcriptional secondary structures of flanking introns and promotes circRNA formation by bringing splice sites of distal exons into proximity of each other ^23,24^. Both back-splicing and linear splicing use fundamental RNA processing components such as the spliceosome machinery and canonical splice sites ^18,25^, which suggests that the two processes occur in a competitive manner. Specific RBPs have been shown to influence circRNA expression in developmental transitions and in various cellular contexts. During epithelial-mesenchymal transition, the splicing factor Quaking regulates the formation of over one-third of abundant circRNAs through intron binding ^26^; in motor neurons, the RBP FUS interacts with the pre-mRNA to control the biogenesis of specific circRNAs ^27^, and the RNA processing factor NOVA2 promotes the biogenesis of numerous circRNAs in mouse cortical neurons ^28^. The mechanisms governing circRNA regulation *in vivo* are yet to be fully elucidated; in particular, the intriguing prevalence of circRNAs in the nervous system indicates the involvement of a general mechanism or effector that shifts the balance between linear and circularizing splicing.

ELAV/Hu proteins are a family of highly conserved RBPs, of which at least one member is expressed in a nervous-system-specific manner and used as marker of neuronal identity across animals ^29–31^. *elav* null mutations are embryonic lethal and ELAV activity is essential in neurogenesis for establishing and maintaining the neuronal RNA transcriptome via alternative splicing and alternative polyadenylation ^32–34^. This regulation occurs co-transcriptionally, with ELAV proteins targeting sequence elements at the 3’ ends or splice sites. ELAV’s global role in RNA processing prompted the hypothesis that the RBP may be involved in co-transcriptional RNA circularization in the nervous system. In this work, we unveil a previously unrecognized role of ELAV proteins as key mediators of circRNA expression in the nervous system, and provide mechanistic insights into circRNA biogenesis.

## Results

### Characterization of the circRNA landscape in Drosophila embryos

In order to identify neuron-enriched circRNAs *in vivo*, we compared transcriptomes in distinct cell populations. We crossed heterozygous *Δelav* or *ΔelavΔfne* mutant parental flies and collected embryonic progeny at two stages of late embryogenesis, 14–16 and 18–20 hours after fertilization. The ELAV paralogue FNE can partially compensate for ELAV functions in later embryonic stages ^32,35^, therefore we used *ΔelavΔfne* mutants for the 18– 20h time point (Fig. 1A and S1A). Embryonic tissues were dissociated into individual cells, fixed, fluorescently labeled and flow-sorted into three distinct populations: wild-type neurons, *Δelav* mutant neurons, and pooled cells from all other embryonic tissues (“non-neurons”). RNA sequencing and gene expression profiling confirmed the purity (Fig. 1B and S1B) and identity (Fig. 1C) of each population.

**Figure 1.**
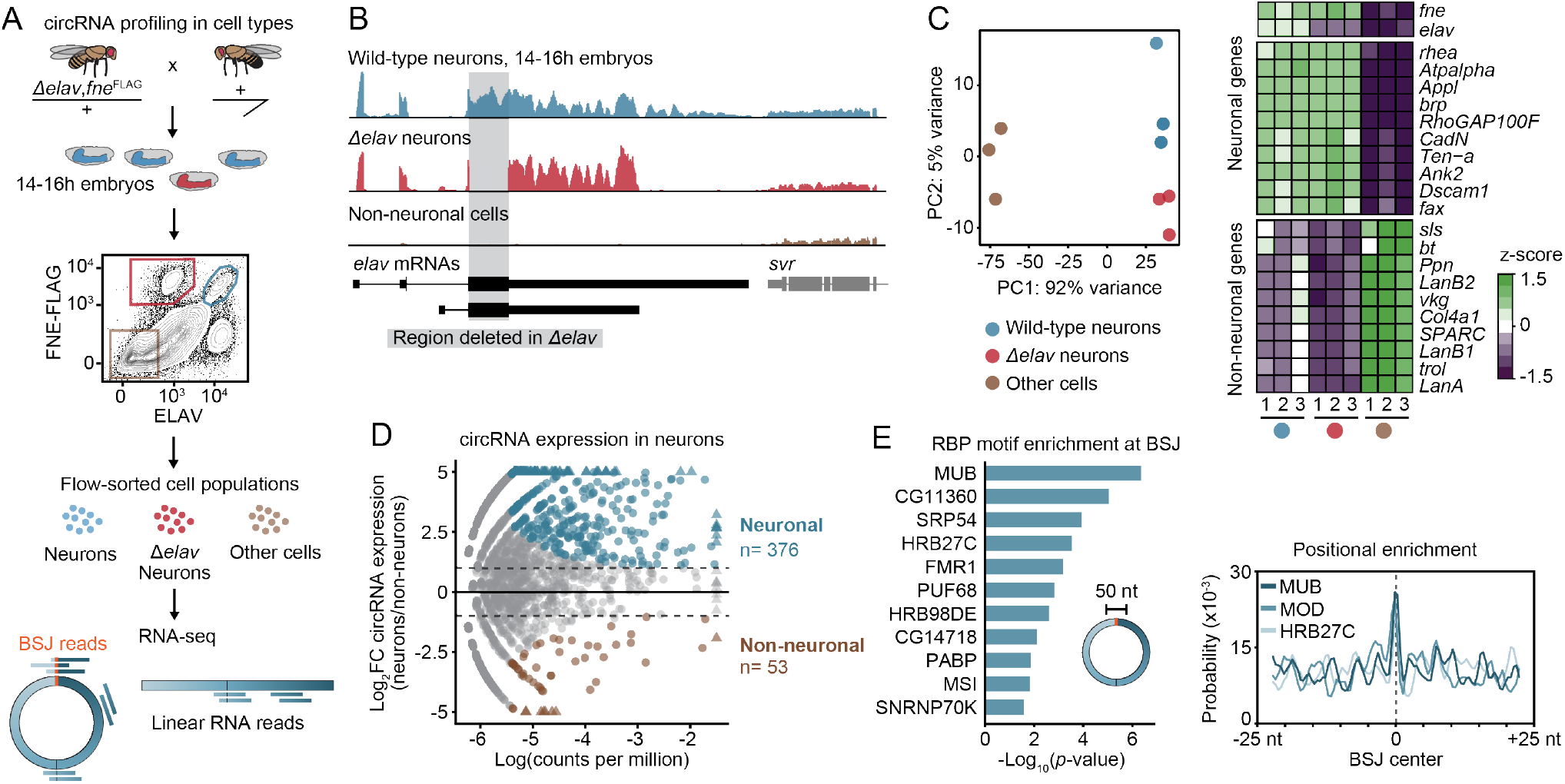
The circRNA landscape in Drosophila embryos. See also Figure S1 and Table S1. (A) Experimental overview: cells from embryonic progeny of *Δelav,fne^FLAG^*heterozygous flies (carrying an *elav* null mutation recombined with a Flag-tagged allele of the neuronal marker FNE) were FACS-sorted into three distinct populations: wild-type neurons (ELAV+, Flag+), *Δelav* mutant neurons (ELAV-, Flag+), and non-neuronal cells (ELAV-, Flag-). circRNA expression was quantified from total RNA-seq data, measuring reads spanning the back-splice junction (BSJ reads) unique to circRNAs. (B) Total RNA-seq tracks at the *elav* locus, in the sorted cell populations. The loss of signal in the *elav* coding region in *Δelav* neurons is highlighted. (C) Principal-component analysis plot of gene expression across the embryonic cell populations. The heatmap on the right indicates the top differentially expressed genes between neuronal and non-neuronal populations. Replicates and identity of cell populations are indicated with colored dots. (D) Differential circRNA expression in neurons compared to non-neuronal populations represented as a function of BSJ counts per million. Significantly enriched (neuronal) circRNAs (p<0.05 and Log(CPM)>-5.4, Log2FC≥1, blue) or depleted (non-neuronal) circRNAs (Log2FC≤-1, brown) are highlighted. (E) Left, RBP binding motifs significantly enriched at the BSJ (±25 nt) of neuronal circRNAs, compared to the entire linearized circRNA sequence. Right, positional enrichment of binding motifs for three neuronal RBPs are indicated from the center of the back-splice junction (right).

To annotate and quantify circRNAs, we counted RNA-seq reads that span back-splice junctions (BSJs) using CIRI2 ^36^, including only circRNAs containing canonical splice signals. We treated bulk tissue samples from adult fly heads and 14–20 h embryos with RNaseR, a treatment that enriches for circRNAs, and compiled a circRNA reference transcriptome by pooling BSJ reads from all sample replicates (Fig. S1C). We subsampled BSJ reads from each dataset and assessed the number of identified circRNAs in each fraction for each cell population. Near-saturation was achieved for circRNAs detected with 5 or more BSJ reads (Fig. S1D); therefore, we set this value as a threshold for stringent circRNA identification and subsequent analyses, in line with previous good practice large benchmarking studies ^37^.

58% of the circRNAs identified (≥5 BSJs) in the sorted cell populations were also present in the RNaseR-treated samples (Fig. S1E, F), validating the accuracy of our annotation and indicating that isolating cell populations helps identify circRNAs that are expressed either lowly or in a tissue-specific manner. Comparing BSJ expression between neuronal and non-neuronal populations with stringent cut-offs, we identified around 400 circRNAs specifically enriched in neurons at either developmental time (hereafter referred to as “neuronal circRNAs”, Fig. 1D and Fig. S1G-J, Table S1), with only 55 circRNAs depleted compared to other embryonic cells (“non-neuronal”). The majority of circRNAs enriched in the neuronal population are transcribed from genes associated with neuronal functions, with a notable enrichment for synaptic signaling and complex behavior (Fig. S1K). This aligns with research in mouse tissues demonstrating that host genes of brain-specific circRNAs are highly enriched for synaptic proteins ^6^, and indicates that circRNAs have a conserved function in the regulation of synaptic processes.

BSJs are unique to circRNAs and may constitute binding sites for RBPs that regulate the circRNA in a manner distinct from that of the linear transcript. When we compared back-splice junction regions of neuronal circRNAs with those of broadly expressed circRNAs, we observed a significant enrichment for neuronal RBP motifs (Fig. S1L, M). Interestingly, when comparing only BSJ regions (BSJ ±25 nucleotides), we found specific neuronal RBPs enriched, including Mushroom-body expressed (MUB), Musashi (MSI) and Fragile X mental retardation protein (FMR1). Some RBPs displayed exquisite positional enrichment at the center of the BSJ, i.e., the precise region discriminating circRNAs from their linear cognate (Fig. 1E). Individual circRNAs have been shown to act independently in cell-type-specific gene expression regulation in different tissues ^16,38–41^ and were proposed to act as tissue-specific RBP and microRNA “sponges” ^7,13,17^. Our results show that circRNAs represent molecular signatures of developing neurons, and suggest they perform cellular functions distinct from those of linear transcripts.

### ELAV mediates neuronal circRNA expression

The RBP ELAV is a key regulator of neuron-specific alternative RNA processing across neuronal cell types, including the generation of neuron-specific (linear) splice isoforms ^32–35,42^. To test a role for ELAV in circRNA expression, we compared the circular transcriptome of wild-type flow-sorted neurons to those of *Δelav* and *ΔelavΔfne* neurons. We found a marked and global decrease in neuronal circRNA expression in absence of ELAV proteins, with over 75% neuronal circRNAs significantly downregulated (Fig. 2A, B and Fig. S2A). circRNA depletion was more pronounced in *ΔelavΔfne* double mutants (Fig. 2C, D and Fig. S2B), showing that FNE rescues circRNA-related functions of ELAV like it does alternative linear RNA processing ^32–34^.

**Figure 2.**
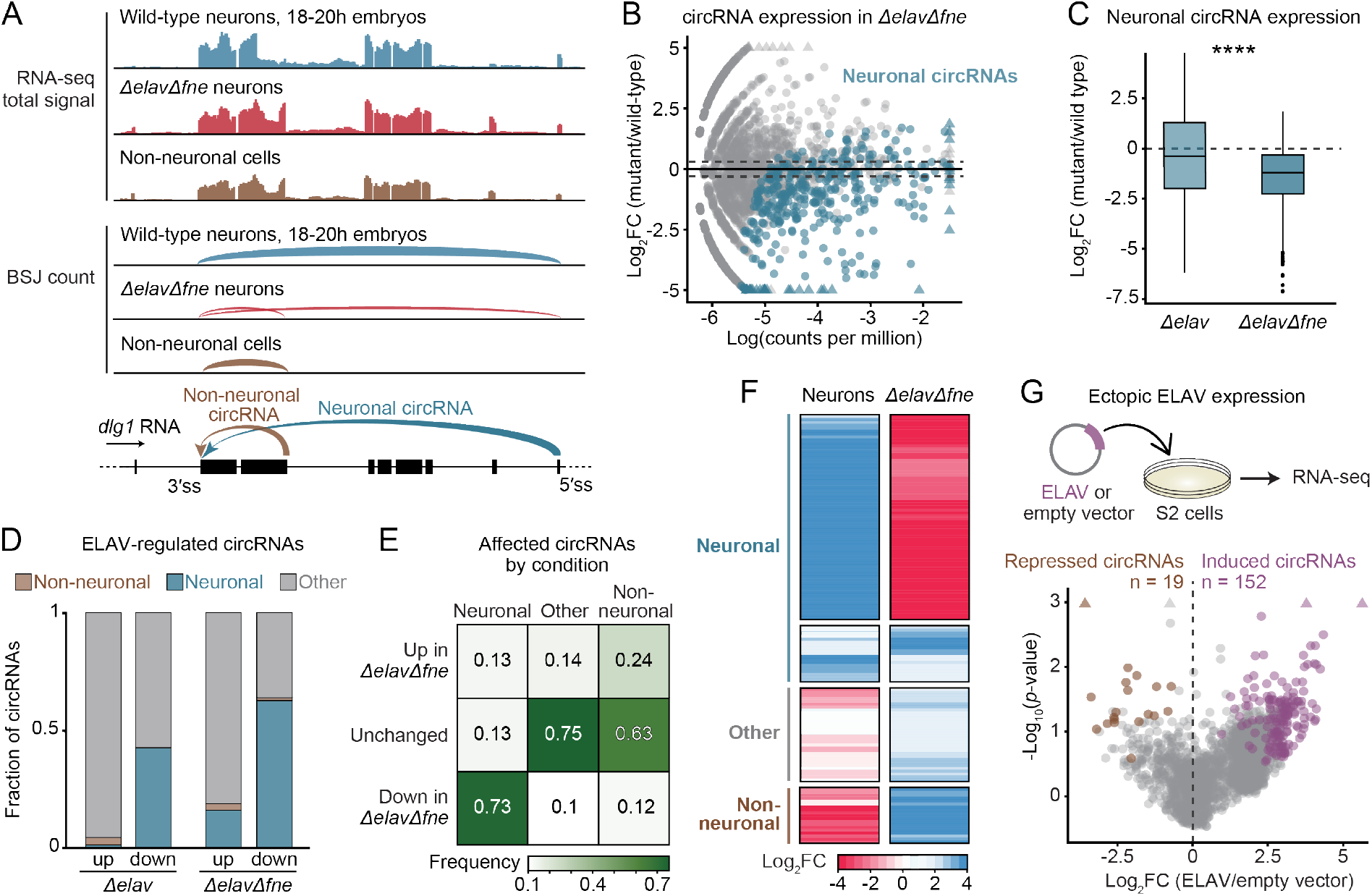
ELAV regulates neuronal circRNA formation. See also Figure S2 and Table S1. (A) Total RNA-seq signal tracks and representation of the BSJ count of a portion of the gene *discs large 1* (*dlg1*), in the sorted cell populations. Arrows in the gene model indicate the back-splicing events that produce the neuronal (blue) or non-neuronal (brown) circRNAs, respectively. (B) Differential circRNA expression in wild-type neurons compared to *ΔelavΔfne* neurons, represented as a function of BSJ counts per million. Highlighted dots represent circRNAs classified as neuronal (Fig. 1D, 376 neuronal circRNAs). The dotted line indicates the abs(log2FC)≥0.3 cutoff. (C) Differential circRNA expression in *Δelav* and *ΔelavΔfne* mutant neurons compared to wild-type neurons. ****p<0.0001 (two-tailed Welch’s t-test). (D) Proportion of circRNAs with neuron-specific expression (neuronal, non-neuronal, other) affected in *Δelav* and in *ΔelavΔfne* neurons. circRNAs were considered significantly affected (up or down) in mutant compared to wild-type neurons if abs(Log2FC)≥0.3 (*ΔelavΔfne*) or abs(Log2FC)≥0.5 (*Δelav*), and Log(CPM)>-5.4. (E) Confusion matrix displaying the fraction of circRNAs with neuron-specific expression that are affected (up, down, unchanged) in *ΔelavΔfne* mutants. (F) Heatmap representing circRNA expression in neurons (differential expression in neurons compared to non-neurons) and in *ΔelavΔfne* (differential expression in mutant compared to wild-type neurons). circRNAs are grouped according to their expression in cell populations (neuronal, non-neuronal, other). Only circRNAs significantly affected in *ΔelavΔfne* are represented. (G) Differentially expressed circRNAs upon ELAV expression in S2 cells compared to an empty vector control. Significantly (p<0.1) upregulated (Log2FC≥0.33, purple) and downregulated (Log2FC≤-0.33, brown) circRNAs are highlighted. Dots are jittered to reduce overlap.

Clustering of circRNA expression in *ΔelavΔfne* neurons identified a distinct pattern: neuronal circRNAs were globally downregulated in *ΔelavΔfne* mutants while non-neuronal or “other” circRNAs were upregulated (Fig. 2E, F), showing that ELAV drives the neuronal identity (“neuronality”) of the circRNA landscape. Additionally, we noted the appearance of circRNAs in *ΔelavΔfne* neurons not detected in any other cell population, “ectopic circRNAs”, most of which were undetectable (BSJ<1) even in RNaseR-treated wild-type tissues (Fig. S2C and Fig. S2D).

Next, we expressed ELAV in Drosophila S2 cells, which are macrophage-like with very low natural ELAV expression. Ectopic ELAV caused an upregulation of many circRNAs, with very few downregulated (Fig 2G), indicating that ELAV constitutes a general, positive regulator of circRNA expression. Thus, it is likely that the pan-neuronal expression of ELAV protein is the main cause for the high circRNA diversity and abundance in neural tissues.

### ELAV binds to nascent RNAs but not mature circles

To test whether ELAV directly binds neuronal circRNAs *in vivo*, we performed ELAV RNA immunoprecipitation with UV-crosslinking followed by RNA sequencing (xRIP-seq) in extract from adult head tissue (Fig. 3A). We ensured identification of high-confidence targets by using a polyclonal antibody directed against native ELAV ^32^, and in an independent experiment, an anti-Flag antibody in head extract of flies in which the endogenous *elav* gene was N-terminally Flag-tagged (*elav^FLAG^*flies). Control xRIP-seq samples were obtained from head extract of untagged flies treated with the anti-Flag antibody. As expected, xRIP-seq recovered almost all transcripts previously identified as ELAV functional targets for AS or APA ^32–35^ (Fig. S3A). However, we found that ELAV does not bind circRNAs: BSJs were strongly underrepresented in xRIP samples compared to input (Fig. 3B, C, and S3B-D). In contrast, xRIP samples were highly enriched in reads originating from genes that host neuronal circRNAs (Fig. 3B, D, and Table S2). In order to discriminate between pre-mRNA and mRNA binding, we performed an analysis based on eisaR ^43^: we compared the enrichment of signals from exon-intron boundaries (pre-mRNA) against that from exon sequences (pre-mRNA and mRNA) for each protein-coding gene. Remarkably, 90% of genes from which neuronal circRNAs originate were enriched at the pre-mRNA level (Fig. 3E and S3E, F). From these data, we conclude that ELAV binds to pre-mRNAs of select genes to promote neuronal circRNA biogenesis. Absence of ELAV binding to closed circles could mean that ELAV binds to intronic sequences that will later be spliced out of the pre-mRNA. Indeed, analysis of ELAV iCLIP data from adult fly heads ^32^ revealed a significant ELAV binding enrichment in introns that flank neuronal circRNAs (Fig. 3F). Together, our data demonstrate that ELAV globally mediates neuronal circRNA expression through binding the pre-mRNA of circRNA host genes.

**Figure 3.**
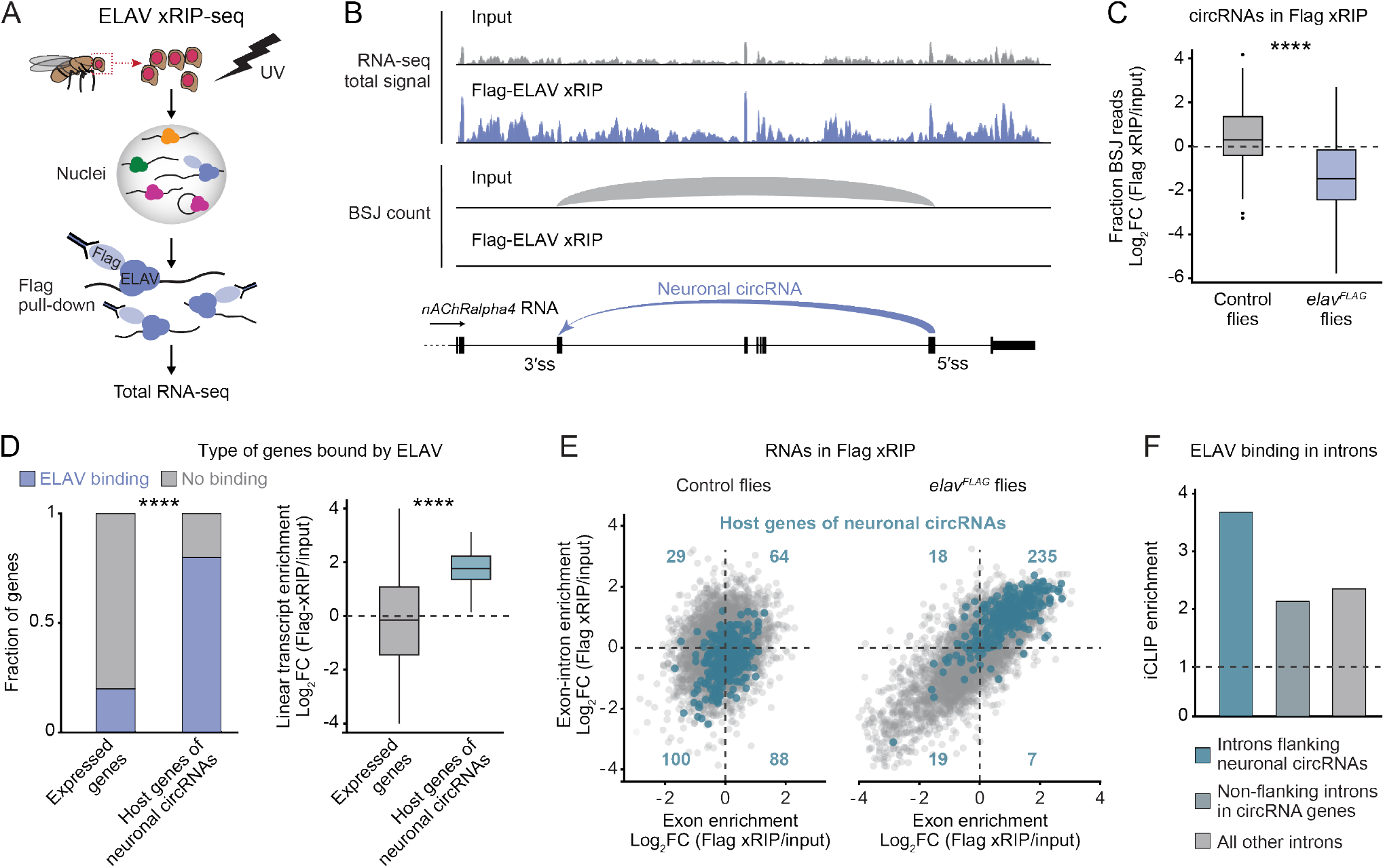
ELAV binds to pre-mRNA of neuronal circRNAs host genes. See also Figure S3 and Table S2. (A) xRIP-seq workflow. UV-cross-linked heads from adult flies underwent nuclear fractionation followed by isolation of protein-RNA complexes and RNA sequencing. In the shown example, Flag-tagged ELAV was captured using anti-Flag beads in flies of the genotype *elav^FLAG^* or *w*^1118^ (control). (B) Total RNA-seq signal tracks and representation of the BSJ count of a portion of the gene *nAChRalpha4* in Flag-ELAV xRIP compared to input. The arrow in the gene model indicates the back-splicing event that produces the neuronal circRNA. (C) ELAV binding to circRNAs, calculated as the ratio of circular to linear transcript expression in Flag-ELAV xRIP compared to input, in *elav^FLAG^* and *w*^1118^ (control) flies. ****p<0.0001 (one-tailed Welch’s t-test). (D) Left, proportion of genes that displayed significant enrichment in Flag-ELAV xRIP-seq compared to input (Log2FC>1 and p<0.01), in host genes of neuronal circRNAs and in all other genes (expressed in input sample). ****p<0.0001 (Pearson’s Chi-squared test). Right, differential linear transcript expression in Flag-ELAV xRIP compared to input, for the same gene groups. ****p<0.0001 (one-tailed Welch’s t-test). (E) Exon-intron split analysis of Flag-ELAV xRIP compared to input. Each dot represents one gene. For each gene, ELAV enrichment at exon-intron boundaries (pre-mRNA) is shown as a function of ELAV enrichment in exons (total transcript). Host genes of neuronal circRNAs are highlighted in blue. Total transcript enrichment and pre-mRNA enrichment were calculated from exon reads and from reads overlapping exon-intron boundaries, respectively. (F) Enrichment of ELAV iCLIP signal in introns. Introns immediately upstream or downstream of neuronal circRNA back-splice sites (flanking neuronal circRNAs) are compared for enrichment against other introns of the same gene (non-flanking) and introns of genes that do not host a neuronal circRNA (all other).

### ELAV binds to RCMs to promote back-splicing

We hypothesize that in neurons, ELAV binds to flanking introns of neuronal circRNAs in the nascent transcripts to promote intron pairing and back-splicing (Fig. 4A). To assess whether the role of ELAV in circRNA biogenesis is linked to its established function in regulating neuronal AS, we quantified (linear) splice junction counts in wild-type and *ΔelavΔfne* neurons (Table S3), thereby identifying neural-specific splice events. We found that the majority of (linear) neuronal splicing events are ELAV-dependent (Fig. 4B, Table S3), consistent with prior studies demonstrating the impact of ELAV proteins in the establishment of neuronal mRNA signatures ^32,34^, and also suggesting that ELAV’s role in AS is more extensive than previously recognized. Notably, most (60%) ELAV-dependent circRNAs are produced from genes with no measurable linear AS, indicating that ELAV can independently regulate linear and circular splicing (Fig 4C). circRNAs are often flanked by long introns ^13^; comparing linear and circular splicing targets of ELAV, we noted a nearly ten-fold difference in median intron length (Fig. 4C). We measured ELAV xRIP signal in introns of genes undergoing either neuronal AS or neuronal back-splicing, and found comparable and high enrichment of ELAV binding in both scenarios (Fig 4D). These data show that the ELAV-dependent generation of neuronal circRNAs constitutes a regulated process, which likely evolved to benefit neuron development and functionality. In contrast, ectopic or ELAV-independent circRNAs did not display such specific regulation (Fig. 4D, see also Figs. 1E, 2D-F and S2D), and may constitute non-functional or even deleterious side-products of linear splicing, as has been proposed to be the case for many mammalian circRNAs ^44^.

**Figure 4.**
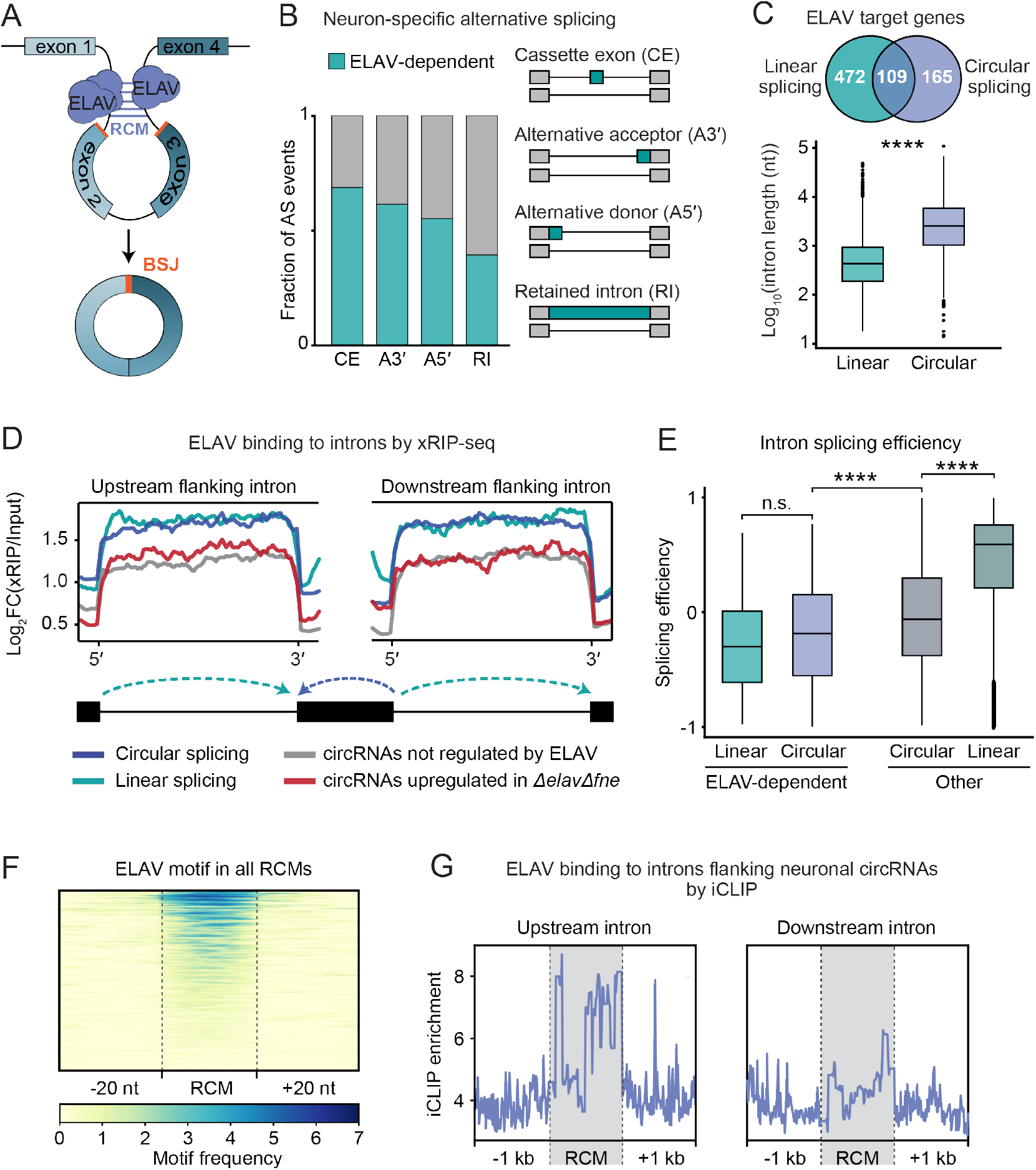
ELAV regulates circRNA production in inefficiently spliced introns. See also Figure S4 and Table S3. (A) Model of ELAV-mediated circRNA biogenesis: ELAV binds to reverse complementary match sequences (RCM) in introns flanking the circRNA back-splice site (BSJ) and inhibits splicing efficiency, resulting in secondary structures that promote neuronal circRNA expression. (B) Proportion of neuronal alternative splicing events (differential splicing events in neurons compared to non-neurons) that are ELAV-dependent (differential in *ΔelavΔfne* compared to wild-type neurons), for the indicated types of splicing. (C) Venn diagram showing the number of genes that undergo ELAV-dependent alternative splicing (linear) and of genes that host an ELAV-dependent circRNA (circular). Below, box plot representing the length of the flanking introns for each type of splicing event. ****p<0.0001 (one-tailed Welch’s t-test). (D) Gene metaplot representing ELAV xRIP-seq enrichment in flanking introns of the indicated types of ELAV-regulated splicing events. Forward and reverse arrows in the gene model represent linear and back-splicing events, respectively. (E) Splicing efficiency of introns undergoing linear splicing or back-splicing, comparing ELAV-regulated AS and circRNA formation to ELAV-independent (other) splicing events. ****p<0.0001 (one-tailed Welch’s t-test). (F) Heatmap showing the number and distribution of ELAV motifs in RCM regions of introns flanking neuronal circRNAs. RCMs of introns that contain at least one ELAV motif (independently of the motif’s position within the intron) are displayed. (G) Profile plot of ELAV iCLIP signal at RCMs and in 1kb surrounding, in introns flanking (upstream and downstream) neuronal circRNAs.

One possible mechanism through which ELAV binding promotes back-splicing is by inhibiting linear splicing of long introns that flank circRNAs. We evaluated co-transcriptional splicing efficiency using nascent RNA-seq data from Drosophila heads ^45^, comparing signal in unspliced introns to that in their respective downstream exons ^46^ (Fig. S4A). Consistent with findings in Drosophila S2 cells ^47^, splicing efficiencies generally decreased with intron length (up until 10 kb; Fig. S4B). Accordingly, the —typically long— introns flanking circRNAs exhibited significantly lower splicing efficiencies than non-circRNA-associated introns (Fig. S4C). Interestingly, ELAV-dependent introns are spliced significantly less efficiently compared to other introns, whether in the context of AS or circRNA formation (Fig. 4E), even within the same gene (Fig. S4C). These observations suggest that ELAV binding reduces splicing efficiency, thereby increasing the window of opportunity for AS and back-splicing. We also hypothesized that ELAV binding may facilitate the formation of secondary structures that promote intron pairing at reverse complementary match sequences (RCMs) (Fig. 4A). RCMs constitute hallmarks of inter-intronic interaction and can predict circRNA formation ^23,24,48^. We identified multiple RCMs in flanking introns of most circRNAs, with a particular prevalence in ELAV-dependent circRNAs (Fig. S4D). We found that RCM regions were highly enriched in ELAV binding motifs (Fig. 4F); ELAV iCLIP from brain tissue showed significant local binding of ELAV at RCMs flanking neuronal circRNAs, with signal particularly strong in the upstream intron, i.e., the intron transcribed first (Fig. 4G). Together, our data show that ELAV regulates back-splicing and neuronal circRNA formation by binding to RCMs in flanking introns.

## Discussion

In this study, we demonstrate that the pan-neuronal protein ELAV is the global mediator of neuronal circRNA synthesis; ELAV’s tissue-specific expression underlies the extraordinary abundance of circRNAs in the nervous system. Mechanistically, ELAV binds to the pre-mRNA of genes that produce neuronal circRNAs, specifically targeting RCM sequences in BSJ-flanking introns. Long introns are associated with circRNA formation in flies (this study; ^13^) and mammals ^49^ as well as with low splicing efficiencies ^50,51^, which supports the notion that many circRNAs constitute functionally neutral or deleterious side-products of splicing. Our data indicate that most neuronal circRNAs do not belong to that category: ELAV-dependent circRNA formation occurs in a regulated and highly specific fashion, with reduced splicing efficiencies in ELAV-bound introns. In our model, ELAV binding to the RCM in the upstream flanking intron represents the crucial step, ensuring retention of that intron and its RCM while the downstream intron is being transcribed, which can take several minutes in long neuronal genes. It is still not clear how ELAV achieves the targeting specificity required to induce circularization at distinct neuronal BSJs. The ELAV binding motif is found broadly distributed across the genome and can by itself not account for the observed positional enrichment at RCMs. This is reminiscent of the high prevalence of Alu elements in the introns of primates, where Alu pairing regulates linear ^52,53^ and circular ^54^ splicing. Even within the nervous system, circRNAs display cell-type-specific expression patterns ^15,55,56^; it is possible that the combination of transcription and RNA processing factor specificities selects and modulates transcript circularization in distinct neurons, perhaps even in distinct synapses.

In fly embryos, at least 75% of neuronal circRNAs depend on ELAV/FNE. The accumulation of circRNAs in the brain is a molecular landmark across animals ^1,12,13^. Considering the similarity, in terms of sequence and expression patterns, of neuronal ELAV-like (nELAV) proteins from flies to humans ^29,57,58^, conserved mechanisms may be at play to regulate intron pairing and neuronal circRNA biogenesis. Moreover, the predominantly cytoplasmically localized human nELAV protein HuD binds to a quarter of all brain-expressed circRNAs as well as many host transcripts ^59^; in context with our findings, this raises the hypothesis that in evolutionarily distant species, nuclear nELAV proteins (such as ELAV) mediate circRNA biogenesis, and that binding of mature circRNAs by cytoplasmic nELAVs (such as HuD) regulates their local expression and function. The exquisite co-regulation of neuronal circRNAs at the levels of biosynthesis (this study, ^28^), subcellular localization, and response to extrinsic signals ^6,12,60^, could indicate that neuron-specific, nELAV-dependent circRNAs may function in concert, for example by binding a common set of RBPs. Our finding that BSJ regions of neuronal circRNAs are enriched for binding motifs for neuronal RBPs are consistent with this possibility. RBPs often localize in synapses, where they regulate the local translation and transport of neuronal transcripts ^61,62^. With their long half-life, unique RBP binding sites, and versatile functionalities, circRNAs possess great potential for modulating gene expression in a cell-specific and even synapse-specific manner.

### Limitations of the Study

Our cell sorting approach aimed at enriching neuronal cell types, providing insights into nervous system specific circRNA expression. It is important to note that circRNAs can exhibit highly cell-type specific expression patterns. This specificity cannot be resolved in our sorting methodology. To address this limitation, future studies using single-cell RNA sequencing technologies are essential. Such approaches would allow for the high-resolution detection of circRNAs within individual brain cells. Additionally, single-cell studies could uncover the dynamic regulation and functional roles of circRNAs in diverse neuronal populations, contributing to a deeper understanding of their involvement in brain development and neurological diseases.

## Acknowledgements

We thank Fernando Mateos for technical help. We are grateful to Alejandro Gomez Auli and Gerhard Mittler at the Proteomics Core, Ulrike Bönisch and the Deep Sequencing Core, and Thomas Manke and the Bioinformatics Core at MPI-IE. We thank Andreas Lingel for his help in designing *elav^RBD^*. We thank Hasan-Can Ozbulut and Nikolaus Rajewsky for helpful discussions, and Anton Heß for critical reading of the manuscript. Stocks obtained from the Bloomington Drosophila Stock Center (NIH P40OD018537) were used in this study. This work was funded by the Max Planck Society, the Deutsche Forschungsgemeinschaft (DFG, German Research Foundation) SFB 1381 (Project-ID 403222702) and under Germany’s Excellence Strategy (CIBSS-EXC-2189-Project-ID 390939984), and the European Research Council (ERC) under the European Union’s Horizon 2020 research and innovation program (grant agreement ERC-2018-STG-803258).

## Author contributions

C.A.-G. and V.H. conceptualized the study. C.A.-G., S.H., S.G., M.S., J.C., and S.T. performed experiments. C.A.-G., S.H., S.G., M.S. and V.H. designed and analyzed experiments. C.A.-G. and V.H. designed computational data analysis. C.A-G. and M.R. performed computational data analysis. C.A.-G. and V.H. prepared the figures. C.A.-G. and V.H. wrote the manuscript with input from all authors. V.H. supervised the study and acquired funding.

## Supplemental Information

**Document S1.** Figures S1–S4.

**Supplemental Tables.** Tables S1-S4. Excel files containing additional data too large to fit in a PDF.

**Table S1. CircRNA identification and quantification in flow-sorted cell populations from Drosophila embryos. Related to Fig. 1 and Fig. 2.** circRNAs identified as differentially expressed in each indicated dataset (neurons vs. non-neurons at 14–16h, neurons vs. non-neurons at 18–20h, *Δelav* neurons vs. wild-type neurons at 14–16h, *ΔelavΔfne* neurons vs. wild-type neurons at 18–20h), with quantification results.

**Table S2. xRIP-seq identification of transcripts directly bound by ELAV. Related to Fig. 3.** Transcripts identified as enriched in each indicated dataset (anti-ELAV antibody xRIP vs. input in wild-type flies, and anti-Flag antibody xRIP vs. input in *elav^FLAG^* flies), with quantification results.

**Table S3. Identification and quantification of alternatively spliced exons in flow-sorted cell populations from Drosophila embryos. Related to Fig. 4.** Exons identified as differentially expressed in each indicated dataset (neurons vs. non-neurons at 18–20h, *ΔelavΔfne* neurons vs. wild-type neurons at 18–20h), with quantification results.

**Table S4. Recombinant DNA and RT-qPCR oligonucleotides used in this study. Related to STAR Methods.**

## STAR Methods

### Contact for reagents and resource sharing

Further information and requests for resources and reagents should be directed to and will be fulfilled by Valérie Hilgers.

### Data and code availability

● All sequencing data generated during this study can be accessed at NCBI Gene Expression Omnibus under the accession number (GSE269179).
● Code used in this study to generate all figures is deposited in https://github.com/hilgers-lab/circles2024 and https://github.com/hilgers-lab/SpliceFlow.
● Any additional information required to reanalyze the data reported in this paper is available from the lead contact upon request.

### Experimental model and genome editing

In this study, all experiments used *Drosophila melanogaster* male embryos. The age of the embryos is specified in hours after egg laying (AEL) when maintained at 25°C. Control flies (*w*^1118^) and GFP-marked balancer chromosomes were sourced from the Bloomington Stock Center (lines 5905, 4559, 6662). Null alleles for *fne*, denoted as *Δfne*, were acquired from M. Soller (Zaharieva et al., 2015)^63^; *Δelav* and *fne^FLAG^* alleles are from (Carrasco et al., 2020)^32^. *elav^FLAG^* and *elav^RBD^* fly lines were generated through CRISPR/Cas9 genome editing, adhering to the methods outlined by Port and Bullock ^64^. Embryo injections were performed by Bestgene, Inc. *elav^FLAG^* expressing an endogenously, N-terminally Flag-HA-tagged ELAV protein, was generated using a guide RNA (GTCTACTCCGCCGCCAGCTC) targeting *elav* and a plasmid comprising 1406 nucleotides upstream of the tag insertion site, the Flag-HA sequence, and 1509 nucleotides downstream the insertion site (sequence in Table S4). To create *elav^RBD^*, a total of ten single-nucleotide mutations in all six RNP motifs of the three ELAV RRMs, previously described to eliminate ELAV RNA-binding (Lisbin, M. et al., 2000), were introduced by injecting the *elav^FLAG^* line with a plasmid containing two guide RNAs (AACCACAGCAGGCGCAGCCC, GTCTACTCCGCCGCCAGCTC) and a 1257-nucleotide gBlock (Integrated DNA Technologies) as a homology donor (sequence in Table S4). The resulting amino acid mutations are the following: RRM1, RNP1: Y205A and F207D. RRM1, RNP2: I152A and N154A. RRM2, RNP1: F294A. RRM2, RNP2: Y251A. RRM3, RNP1: Y445A and F447D. RRM3, RNP2: F405A and Y407A.

### Genetic strategy to sort wild-type and Δelav mutant neurons (14–16h dataset)

We used the progeny of *Δelav, fne^FLAG^/FM7* and FM7/Y flies that contain the *elav* null allele *Δelav* ^32^ recombined with an endogenously V5-flagged *fne (fne^FLAG^*) ^65^. Flag was used as a marker of neuronal cells and ELAV protein was used as a marker of wild-type neurons, which allowed to distinguish *Δelav* mutant neurons (Flag+, ELAV-), wild-type neurons (Flag+, ELAV+), and non-neuronal cells (Flag-, ELAV-).

### Genetic strategy to sort wild-type and *ΔelavΔfne* mutant neurons (18–20h dataset)

Flies described as *ΔelavΔfne* flies are progeny of *elav^RBD^,Δfne*/FM7 crossed with FM7/Y males. As a result, the RNA-binding dead ELAV protein can be discriminated from the wild-type ELAV by Flag, and never co-expresses with FNE. In heterozygous flies, the *elav^RBD^* allele is robustly suppressed (through a mechanism we have not characterized), leading to the absence of detectable *elav^RBD^* protein (Figure S1A). This suppression allows for the positive identification of neuronal populations that lack the ELAV^RBD^ protein using Flag as a marker. Wild-type neurons (Flag- and ELAV+) were distinguished from mutant neurons (*elav^RBD^,Δfne*: Flag+, ELAV+), and non-neuronal cells were identified as (Flag-, ELAV-).

### RNaseR treatment

For head samples, 3-day-old w1118 flies were collected, flash-frozen in liquid nitrogen, decapitated by shaking and heads were manually collected with a thin brush. For embryo samples, eggs from *w*^1118^ flies were collected for two hours on agar plates and aged for 14h (14–16h AEL embryos) at 25°C. In both cases, samples were homogenized in QIAzol Lysis Reagent (QIAGEN 79306) for RNA extraction. 2µg RNA per sample was treated with 10 U RNAseR (Applied Biological Materials, E049) in 20 µL for 20 min at 37°C. For non-treated control samples, the enzyme was omitted. After addition of spike-in mouse embryonic stem cell RNA at 0.33 ng/µL final concentration, RNA was extracted with Trizol LS (Ambion 10296028) and glycogen (Invitrogen AM9515) according to the manufacturer’s protocol. RNAs were reverse transcribed using Maxima RT (Thermo Scientific EP0741) with random hexamers (Jena Bioscience 94824), according to the manufacturer’s protocol.

### Total RNA sequencing library preparation

RNA integrity was analyzed using a 2100 Bioanalyzer (Agilent Technologies). Libraries for total RNA-seq were prepared with 100 ng of total RNA using TruSeq Stranded total RNA Library Prep (Gold) (Illumina 20020599) according to the manufacturer’s instructions. Paired-end sequencing was performed using the NovaSeq6000 platform (Illumina) and 101-bp reads.

### Fluorescence-activated cell sorting in Drosophila embryos

Drosophila embryos were collected at the appropriate times after egg laying. Sorting of embryo populations was done based on (McCorkindale et al., 2019)^66^, with modifications. All steps until FACS were carried on at 4°C, and all washes used cold, RNAse-free PBS (Alfa Aesar, J62851) with RNase inhibitor (1:250) (Ribolock, ThermoFischer) and 1000g for 2 min to preserve RNAs from degradation. Embryos were dounced in RNAse-free PBS with RNase inhibitor (1:250), and cell debris were filtered out by filtration through a 40µm strainer at 400g. Cells were washed twice, and stained with Zombie Aqua (BioLegend, 423101) for live/dead separation for 15 min in the dark, in PBS with 1:100 RNase inhibitor. Cells were washed twice and then fixed with 4% formaldehyde (ThermoFischer scientific, 28908) in PBS for 15 min in the dark. Fixation was stopped with incubation for 3 min with a quenching buffer (750 mM TrisHCl pH 7.5 in DEPC-PBS, 1:100 RNase inhibitor) and cells were washed again twice. Cells were incubated with the following primary antibodies: polyclonal anti-ELAV 1:500 (rabbit, generated in Carrasco et al., 2020)^32^ and mouse-anti-Flag M2 (Sigma, F1804) at 1:500 for 45 min in the dark, with agitation, at 4 °C in staining buffer (1% BSA (w/v) (Sigma, 6917), 0.1% saponin, 1:200 RiboLock). Cells were washed twice in 0.2% BSA, 0.1% saponin in PBS, 1:100 RiboLock and incubated with fluorescently conjugated antibodies at 1:500 (14–16h embryos) or 1:1000 (18–20h embryos) for 45 min in the dark, with agitation, at 4 °C in staining buffer. Sorting used 488 anti-mouse (Abcam) and 555 anti-rabbit (Invitrogen) in *Δelav* embryos; and 488 anti-rabbit (Life Technologies) and 555 anti-mouse (Abcam) in *ΔelavΔfne*. Cells were then washed twice, resuspended in sorting buffer (0.5% BSA, 2mM EDTA, 1:500 RiboLock), filtered through a 40µm nylon mesh, sorted using a FACSymphony (Becton Dickinson), and collected in cold sorting buffer. Cells were pelleted and RNA-protein complexes were reverse-cross-linked with proteinase K (Ambion AM2546) in 10mM Tris, pH8, 10mM NaCl, 1mM EDTA, 1:100 proteinase K, 1:100 RiboLock), for 20 min at 50° C. RNA was extracted in 3:1 Trizol LS (Ambion) according to the manufacturer’s protocol.

### Cell transfection

S2 cells were transfected in six-well plates using Effectene (Qiagen) with 300 ng of the λN expression plasmid: λN empty plasmid or λN-ELAV from Hilgers et al., 2012 ^67^. Transfections were conducted in three replicates. Cells were harvested 48 hours after transfection by centrifugation and lysed in QIAzol Lysis Reagent (QIAGEN 79306) for RNA extraction.

### circRNA validation by qPCR

300 ng of total RNA was used per replicate. RNA was reverse transcribed using Maxima RT (Thermo Scientific EP0741) with random hexamers (Jena Bioscience 94824) according to the manufacturer’s protocol. RT-qPCR was performed in a LightCycler 480 II instrument using FastStart SYBR Green Master (Roche). Divergent primers were designed to amplify only circular RNAs from the target genes. RT-qPCR primer sequences are listed in Table S4.

### Cross-linking RNA-Immunopurification and sequencing (xRIP-seq)

Drosophila heads were separated from bodies by shaking in liquid nitrogen, and ground into powder in liquid nitrogen. For one sample, 100 mg head powder was subjected to UV cross-linking (UV-C, 254 nm) through six rounds at 300 mJ/cm² in a BLX-312 crosslinker (Bio-Link). All subsequent steps were performed at 4°C. Nuclear extracts were obtained by homogenizing the UV-cross-linked head powder in 1 mL homogenization buffer (0.2% Triton X, 10% sucrose, 0.5 mM DTT, double the standard concentration of protein inhibitors (Roche, 11873580001), and 1:500 RiboLock RNase inhibitor (Thermo Fisher Scientific, EO0384), using ten strokes with a loose pestle. For each replicate, three samples (3mL of homogenate) were pooled, homogenate was filtered using a 40µm strainer and centrifuged at 200 g for 3 min. Supernatants were transferred to 15-mL Protein LoBind (Eppendorf, 0030122216) tubes and centrifuged at 800 g for 10 min to pellet nuclei. Pellets were resuspended and centrifuged at 600 g for 10 min to pellet nuclei in 6 mL homogenization buffer excluding Triton X and centrifuged at 800 g for 10 min to pellet nuclei. This pellet was then resuspended in Lysis buffer (50 mM Tris (pH 7), 500 mM LiCl, 10 mM EDTA, 5 mM DTT, 2% lithium dodecyl sulfate (LDS), 0.5% SDS, and 0.5% sodium deoxycholate) and rotated for 10 min. The lysate was centrifuged at 15,000 g for 10 min, and the supernatant transferred to a new tube. After another centrifugation under the same conditions, ionic detergents were neutralized with 1% NP-40. For the Flag-ELAV experiment, the clarified lysate (750 µL; referred to as ‘input’) was incubated with 40 µL anti-Flag M2 magnetic beads (Invitrogen, M8823) for 1.5 hours. The beads were rinsed with lysis buffer (0.1% SDS and 0.1% Na-deoxycholate), washed three times for 5 min with lysis buffer (0.1% SDS, 0.1% Na-deoxycholate, and 1% Triton X-100), and washed twice with lithium chloride buffer (350 mM LiCl, 50 mM Hepes-KOH (pH 7.5), 1 mM EDTA, 1% NP-40, and 0.7% Na-deoxycholate) for 5 min. Protein-RNA complexes were eluted with 120 µL of elution buffer (10 mM Tris-HCl (pH 7.4), 150 mM NaCl, 0.1% Triton X-100, without protease inhibitors) containing Flag peptide (0.2 mg/ml; Sigma-Aldrich, F3290) for 1 hour. The eluates and corresponding inputs underwent treatment with proteinase K (Ambion, AM2546) for 30 min at 50°C and 1100 rpm. For the endogenous ELAV IP experiment, input was incubated with 40uL of the conjugated Dynabeads protein A-antibody complex with anti-ELAV antibody (rabbit, generated in Carrasco et al., 2020)^32^ for 1 hour. Protein-RNA complexes were eluted in 120uL of 1x Proteinase K Buffer (Tris HCl pH 7.5, NaCl EDTA, 6 uL 10% SDS, 1.5 uL Proteinase K (20 mg/mL) and 1 uL Ribolock). RNA was then purified using TRIzol LS Reagent (Ambion, 10296028) following the manufacturer’s protocol. Equal volumes of total RNA from both input and IP samples were used to prepare libraries for total RNA-seq.

### mRNA sequencing data processing

Sequencing data were processed using the RNA-seq module from snakePipes v2.4.3 ^68^, adding flags for --trim, -m “alignment-free,alignment”. Reads were mapped to the *Drosophila melanogaster* reference genome (Ensembl assembly release dm6), and the transcriptome reference annotation release-96 using STAR ^69^. Quality control of RNA-seq reads was done using FASTQC ^70^. To compare gene expression estimates across cell populations, a variance stabilizing transformation (VST) was applied using the DESeq ^71^ function vst() on raw gene counts data from the different samples. The transformed data was used to compute a PCA using the DESeq2 function plotPCA() with standard parameters. Differential gene expression analysis was performed using DESeq2 filtering for baseMean>=4, logFC>1 and padj<0.01 to determine neuronal enriched genes.

### Generation of the circRNA reference database

The circRNA reference database was constructed from a compilation of libraries from RNaseR-treated RNA samples (from heads and embryos) and from RNA samples obtained from flow-sorted embryonic cell populations. For each group, replicates were pooled. Libraries were aligned using Burrows-Wheeler Alignment tool with the parameter “-T 19 “ ^72^, and circRNAs identified using CIRI2 under default settings as recommended by the tool’s developers ^36^, based on the dm6 genome and Ensembl 96 annotations. This process involved filtering and consolidating the identified circRNA entities. Filtering criteria included a minimum of five back-splice junctions (BSJ) per circRNA per cell population, with mutant-specific circRNAs identified by the presence of more than five BSJs across the combined single and double mutant libraries. This consolidated collection of circRNAs formed the reference database that was used in subsequent analyses.

### Saturation analysis (sub-sampling)

Saturation analysis was performed by pooling all replicates from one sample group, and randomly sampling different fractions from 1% to 100% from the raw read files using seqtkV1.2-r94 ^73^. Then, the CIRI2 ^36^ pipeline was applied to each of the sample fractions. Results were summarized as a fraction of recovered compared to the full set.

### Differential circRNA expression

We computed differential circRNA expression by executing the CIRIquant ^74^ workflow and using the circRNA reference database. CIRIquant was run with circRNA in format “ *--ciri --library-type 2* “and using script *prepDE.py* to handle replicates to compute the differential status for each circRNA. After the circRNA quantification with CIRIquant, circRNAs were classified based on their expression in a given embryonic cell population compared to another cell population. We constructed four groups, two for the neuronal comparison (neuronal, non-neuronal) and two for the mutant comparison (ELAV-down, ELAV-upregulated). For classification, we require that a circRNA comply with edgeR parameters of logCPM>-5, p-value<0.1 and |LogFC|>0.

### Motif enrichment at BSJs

RBP enrichment in BSJs of neuronal circRNAs was performed by generating a sequence of 50 nt flanking the BSJ site (±25nt) using the BSgenome.Dmelanogaster.UCSC.dm6 reference genome package in R. The FASTA files were submitted to the MEME suite server and the AME ^75^ program was used to calculate enrichment over the sequences. CentriMO ^76^ was used for assessing positional enrichment. For the comparisons, circRNAs that were neither enriched nor depleted in neurons were used as control sequences.

### Identification of ELAV target genes by xRIP-seq

To identify ELAV target genes, we used the DESeq2 pipeline DESeq ^71^ and assessed read counts in xRIP compared to input. Prior to applying DESeq2, we filtered for genes with more than 10 read counts across all samples and replicates. For xRIP-seq performed with the anti-ELAV antibody in wild-type fly heads, DESeq2 was executed with default parameters using a simplified design: “design = ∼type”, where ‘type’ represents the comparison between input and IP. For xRIP-seq performed with the anti-Flag antibody in *elav^FLAG^* flies, we used a likelihood ratio test with the following parameters: “design = ∼type + condition + condition:type and DESeq(test = “LRT”, reduced = ∼type + condition)”. In this context, ‘condition’ accounts for the confounding factor of non-specific binding to the anti-Flag antibody, which we controlled for by performing Flag IP in untagged *w*^1118^ flies. ELAV target genes for each xRIP experiment were selected with the criteria p.adj<0.05 and log2FoldChange>1.

### Exon-intron split analysis (EISA) on xRIP-seq data

We counted reads specifically spanning exon-exon (EEJ) junctions and exon-intron boundaries (EIB). EIB reads were defined as reads that do not span exon-exon junctions and fully match the reference genome, indicated by having only ‘M’ (match or mismatch) in their CIGAR string from STAR mapping. These reads are assigned to regions that overlap both exons and introns. EEJ and EIB reads were counted using featureCounts ^77^ with the parameters: “featureCounts -p -B -C -t ‘gene’ -g ‘gene_id’ -f -s 2 -J “. EISA analysis was performed using the R package eisaR with parameters from Gaidatzis et al., 2015 ^43^: “runEISA(modelSamples = FALSE, geneSelection = ‘filterByExpr’, statFramework=’QLF’, effects=’predFC’, pscnt=2, sizeFactor = ‘individual’, recalcNormFactAfterFilt = TRUE, recalcLibSizeAfterFilt = FALSE) “. In this analysis, read counts from EEJs were used for mRNA counts and reads of EIBs for intron counts (pre-mRNA). Genes with fewer than 10 EEJ counts on average across replicates were discarded.

### ELAV iCLIP data processing

Previously published ELAV iCLIP data was processed as described in Carrasco et al. 2020. In brief, we used iCount ^78^ to align iCLIP reads and call significant cross-link regions. Intronic cross-link sites were called using iCount and the exonic signal of isoform-specific exons was removed before computing the mean coverage per region across replicates. These signal tracks represent highly confident intronic cross-link sites. For the analysis of ELAV iCLIP signals in flanking introns, we first constructed an intron database to identify sets of introns flanking circRNAs and differentially expressed exons. This enabled us to compute signal enrichment in distinct groups. To create the intron database, we selected protein-coding genes, excluded first/last exons from alternative transcription start sites (TSSs) and alternative transcription end sites (TESs), and discarded non expressed exons. The remaining exons were then subtracted from gene coordinates, resulting in the intron database. We excluded introns smaller than 10 nucleotides. To compute iCLIP enrichment, we selected the following features: i) Upstream and downstream introns immediately flanking a circRNA, ii) Non-flanking introns, iii) All introns of the gene set. To assign introns to circRNAs, we collapsed overlapping circRNAs whose respective 5’ and 3’ ends lie within 10 nt of each other. Introns were assigned to a feature if they were within a 10-nt window. For each set of introns, we assessed enrichment by computing the fraction of iCLIP signal in introns normalized by intron length. The iCLIP profile represents the piled-up count of cross-link sites within normalized intron sizes.

### Detection of differential alternative splicing events

To identify alternative splicing events, rMATS ^79^ was performed using the snakePipes mRNA-seq pipeline, adding the flag “--rMATS”. Splicing events were classified as neuronal if, comparing neuronal and non-neuronal cell populations, the following applied: FDR<0.01 and abs(IncLevelDifference)>1. Parameters for the definition of ELAV-dependent splicing were FDR<0.01 and abs(IncLevelDifference)>0.1.

### Calculation of splicing efficiencies from nascent RNA-seq data

To calculate splicing efficiencies, we used previously published nascent RNA-seq data and a modified quantification method ^46^. First, we generated a reference annotation by selecting the last nucleotides of each intron, i.e., the region covering the last (3’) 30% of each intron’s length. For the downstream exon, the quantification used the nucleotides constituting the first (5’) 30% of the exon’s length. Reads aligned to either the intron or the exon were counted using the *summarizeOverlaps* function from the GenomicAlignments package ^80^. These reads were then normalized using DEXseq ^81^. To quantify splicing efficiencies, we calculated the ratio of reads in introns to reads in exons and subtracted this value from 1. Values closer to 1 indicate a higher splicing efficiency (“faster splicing”), as they represent a lower amount of intron reads relative to exon reads. Values closer to 0 indicate lower splicing efficiencies (“slower splicing”). The methodology and code for this analysis are available at: https://github.com/hilgers-lab/SpliceFlow.

### Prediction of RCMS within flanking intron sequences

To identify RCMs, we used the method described in Ivanov et al., 2015 ^24^. The procedure involves a BLAST alignment for each pair of introns flanking a circRNA to identify all the potential candidates. The parameters we used for the alignments were “-strand minus -word_size 7 -outfmt 0”. RCMs of more than 20 nt in length were used for downstream analysis.

