## Supplemental Figures and figure legends for "ELAV mediates circular RNA biogenesis in neurons"

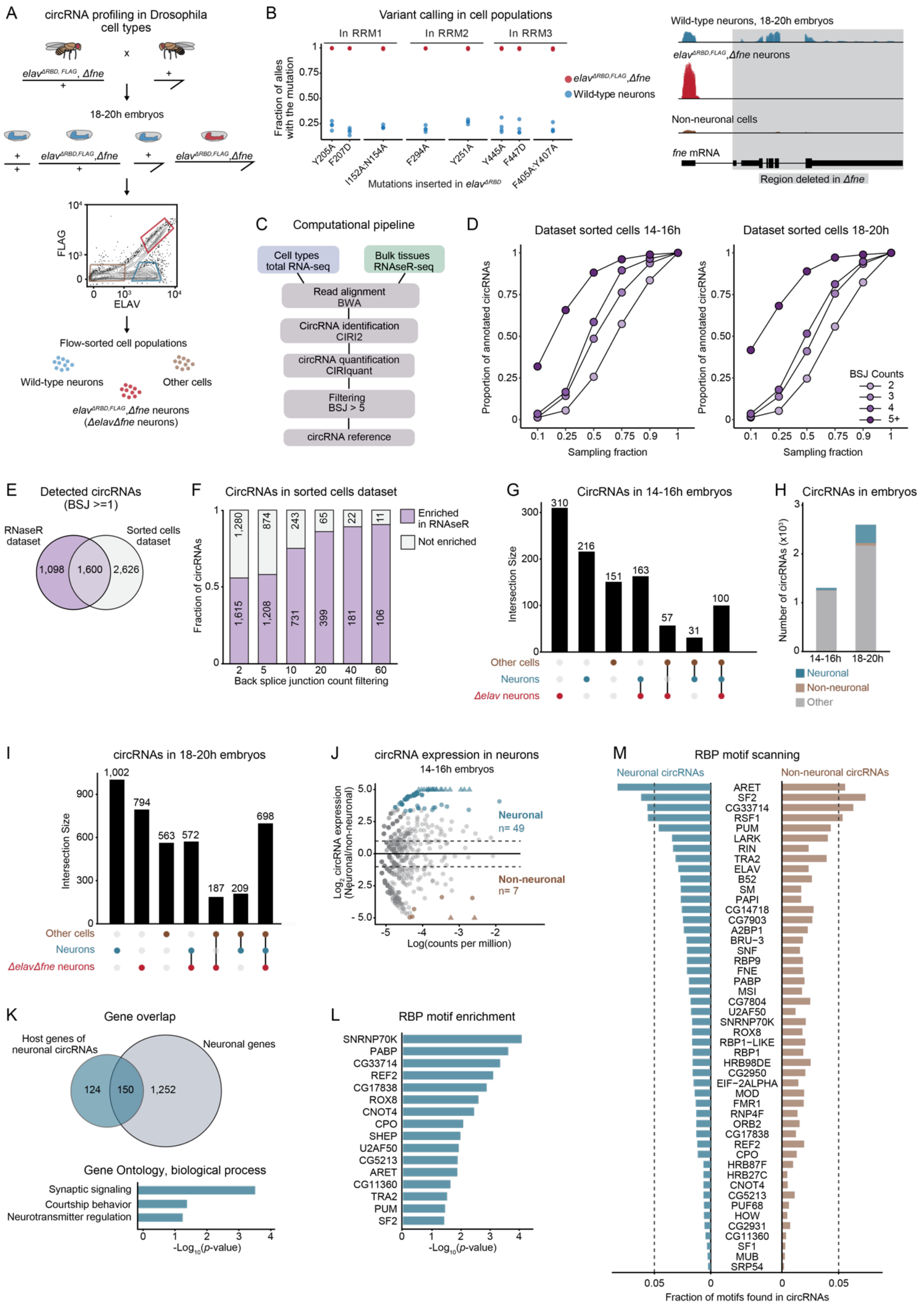


**Figure S1. Landscape of neuronal circRNAs in Drosophila embryos. Related to Figure 1.**

(A) Experimental overview: cells from embryonic progeny of *elav*$^{RBD,FLAG}$*,Δfne* heterozygous flies (carrying a Flag-tagged *elav* RNA-binding dead allele recombined with an *fne* null mutation) were FACS-sorted into three distinct populations: wild-type neurons (ELAV+, Flag-), *ΔelavΔfne* mutant neurons (ELAV+, Flag+), and non-neuronal cells (ELAV-, Flag-). Total RNA-seq was performed on these cell populations in biological replicates. circRNA expression was quantified from total RNA-seq data, measuring reads spanning the back-splice junction unique to circRNAs.

(B) Validation of loss-of-function mutation of *elav* (left) and *fne* (right) in the *ΔelavΔfne* mutant neuron population. Left, variant calling in total RNA-seq data in the indicated cells. Shown is the fraction of reads carrying each indicated mutation, compared to the total number of reads overlapping the mutated region. Each dot represents one sample replicate. Right, total RNA-seq tracks at the *fne* locus in the sorted cell populations. The loss of *fne* signal in *ΔelavΔfne* mutant neurons (downstream of the first exon) is highlighted.

(C) Computational pipeline to build a circRNA reference annotation. For each dataset, reads overlapping back-splice junctions (BSJs) were used to annotate circRNAs using the CIRI2 package (Gao et al., 2020) and quantified using CIRIquant (Zhang et al., 2020). Only circRNAs with ≥5 BSJ counts were considered for the reference annotation.

(D) Saturation analysis of circRNA detection in the indicated RNA-seq datasets, grouped by their expression in BSJ counts. Reads were randomly sampled in the indicated fractions and the circRNA annotation pipeline (omitting the ≥5 BSJ filtering step) was performed in each fraction.

(E) Venn diagram showing the number of circRNAs detected in the indicated RNA-seq datasets before BSJ filtering. Data from replicates and from cell populations obtained from the same embryo time point were pooled.

(F) Proportion of circRNAs enriched in the RNase R dataset, grouped by the number of BSJ counts used for filtering. BSJ≥5 was used as a cutoff for subsequent analyses.

(G) Overlap of circRNAs detected across cell populations sorted from 14–16h embryos.

(H) Number of circRNAs with neuron-specific expression (neuronal, non-neuronal, other) in cell populations sorted from 14–16h and 18–20h embryos. circRNAs were considered significantly enriched in neurons compared to non-neuronal cells (neuronal) if $p<0.05$, Log(CPM)>-5.4 and Log2FC≥1. circRNAs were considered significantly depleted (non-neuronal) if $p<0.05$, Log(CPM)>-5.4 and Log2FC≤-1.

(I) Overlap of circRNAs detected across cell populations sorted from 18–20h embryos.

(J) Differential circRNA expression in neurons compared to non-neuronal populations represented as a function of BSJ counts per million. Significantly enriched (neuronal) circRNAs ($p<0.05$, Log(CPM)>-5.4, Log2FC≥1, blue) and depleted (non-neuronal) circRNAs ($p<0.05$, Log(CPM)>-5.4 and Log2FC≤-1, brown) are highlighted.

(K) Top, Venn diagram showing the number of genes that are host genes of neuronal circRNAs, and of neuronal genes. Neuronal genes are genes whose differential transcript expression was significantly higher ($p<0.01$, Log2FC≥2) in neurons compared to non-neuronal cells in 18–20h embryonic cell populations. Bottom, Gene Ontology classification (GO terms, biological processes) of host genes of neuronal circRNAs. 203 host genes produce 1 circRNA; 71 host genes produce 2 or more circRNAs; in total, 274 host genes produce 396 circRNAs.

(L) RBP motifs significantly enriched at the BSJ (±25 nt) of neuronal circRNAs compared to other expressed circRNAs in cell populations sorted from 18–20h embryos.

(M) RBP motif scanning at the BSJ (±25 nt) of circRNAs, comparing neuronal and non-neuronal circRNAs (in cell populations sorted from 18–20h embryos).

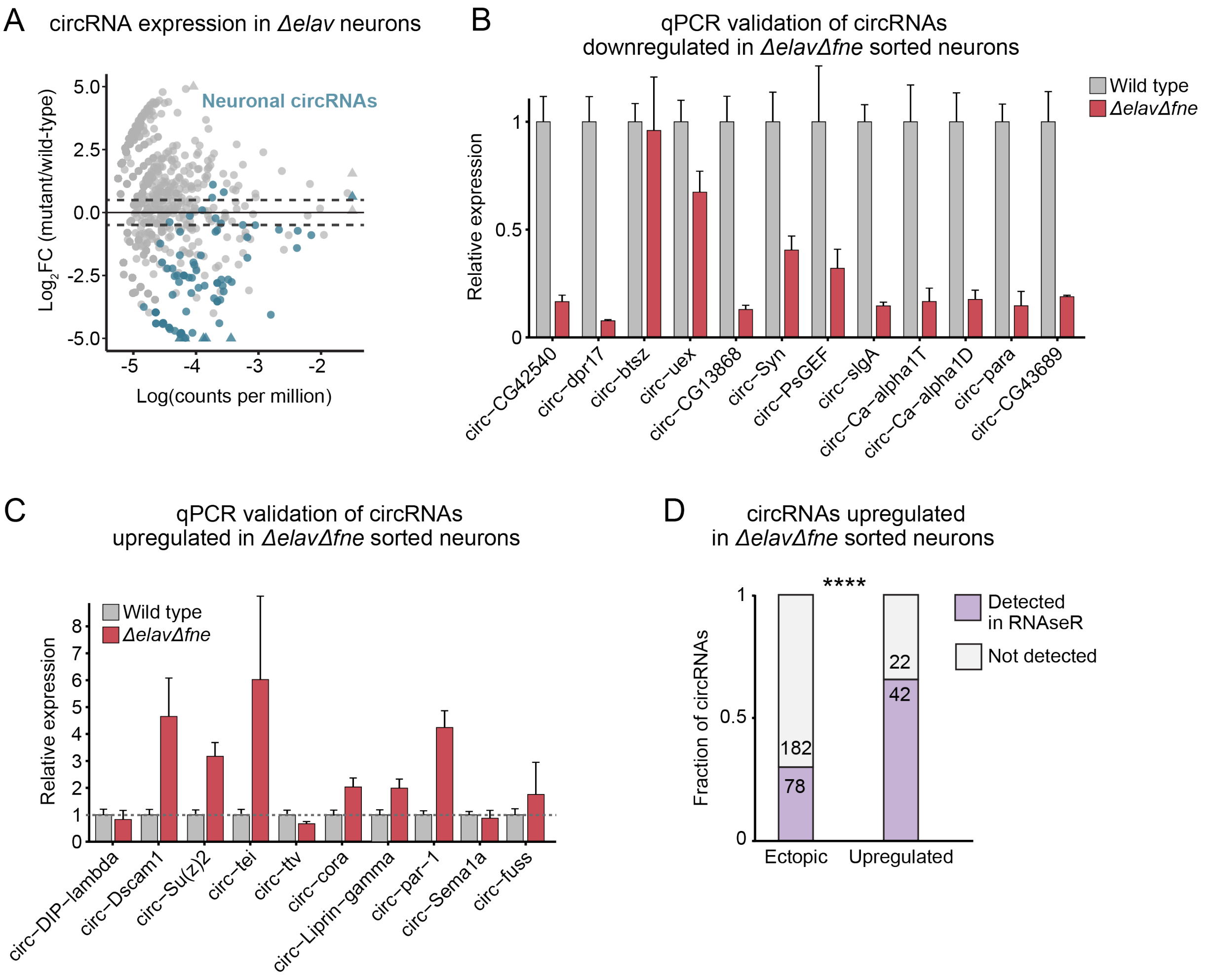


**Figure S2. ELAV regulates circRNA biogenesis. Related to Figure 2.**

(A) Differential circRNA expression in wild-type neurons compared to *Δelav* neurons, represented as a function of BSJ counts per million. Highlighted dots represent circRNAs classified as neuronal in 14–16h embryos (Fig. S1J, 49 neuronal circRNAs). The dotted line indicates the abs(log2FC)≥0.5 cutoff used to classify a circRNA as affected in *Δelav*.

(B, C) RT-qPCR validation and quantification of circRNAs downregulated (B) or upregulated (C) in *ΔelavΔfne* mutant embryos. circRNA levels were normalized to *Rpl32* (*rp49*) mRNA and levels in control flies (wild type) were set to the value 1. Error bars represent mean ±SD of three biological replicates for each genotype. circRNAs were quantified using divergent, BSJ-spanning primers that only detect circular and not linear transcripts of the indicated genes.

(D) Proportion of circRNAs that were upregulated in *ΔelavΔfne* neurons compared to wild-type neurons (ectopic or upregulated) that were detected (≥1 BSJ) in the RNaseR dataset. "Ectopic" denotes circRNAs detected exclusively in the *Δelav* or *ΔelavΔfne* cell population, and not detected in other cell types. "Upregulated" denotes circRNAs significantly upregulated in *ΔelavΔfne* neurons compared to wild-type neurons in cell populations sorted from 18–20h embryos. ****$p<0.0001$ (Pearson's Chi-squared test).

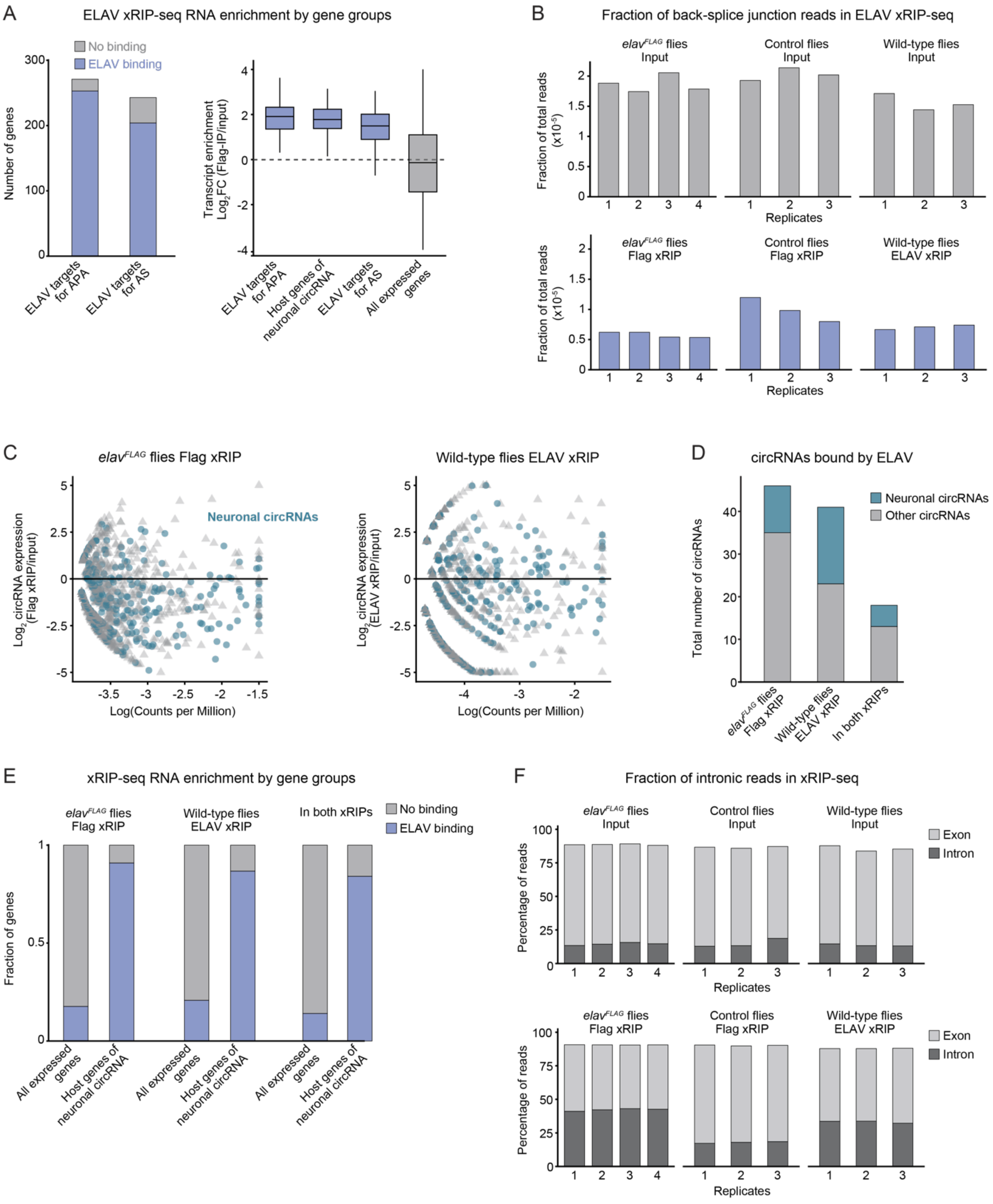


**Figure S3. ELAV binds pre-mRNA of circRNA host genes. Related to Figure 3.**
(A) Left, proportion of genes in each group that displayed significant transcript enrichment in Flag-ELAV xRIP-seq compared to input. “ELAV targets for APA" denotes genes identified in (Carrasco et al., 2020) as genes that undergo ELAV-dependent neuronal alternative polyadenylation. “ELAV targets for AS” denotes genes identified in (Carrasco et al., 2022) and in this study (see Fig. S4) as genes that undergo ELAV-dependent neuronal alternative splicing. Right, ELAV binding to transcripts of the indicated gene groups, calculated as the differential transcript expression in Flag-ELAV xRIP compared to input in *elav*$^{FLAG}$ flies. Genes were considered significantly bound if Log2FC>1 and p<0.01. Genes were considered expressed if they were detected in xRIP-seq and input samples at baseMean>4.

(B) ELAV binding to circRNAs, represented as fraction of total reads that overlap BSJs across replicates, in input (top) and xRIP-seq (bottom) samples in the indicated ELAV pull-down experiments (Flag IP in *elav*$^{FLAG}$ flies, ELAV IP in wild-type flies) and the control experiment (Flag IP in wild-type flies).
(C) Total number of detected circRNAs across samples and replicates of the RIP-seq experiment.
(D) Left, differential circRNA expression in Flag-ELAV xRIP-seq (left) and ELAV xRIP-seq (right) compared to respective inputs, represented as a function of BSJ counts per million. Highlighted dots represent circRNAs classified as neuronal in 18–20h embryonic cell populations. Right, proportion and number of neuronal circRNAs among circRNAs that displayed significant enrichment in Flag-ELAV xRIP, ELAV xRIP, and both xRIPs. circRNAs were considered significantly bound by ELAV if BSJ expression was significantly enriched in xRIP compared to input (Log2FC>0.5 and p<0.1).
(E) Proportion of genes in each gene group that displayed significant transcript enrichment in Flag-ELAV xRIP-seq compared to input in *elav*$^{FLAG}$ flies (Log2FC>1 and p<0.01).
(F) ELAV binding to transcript regions, represented as fraction of total reads that overlap intronic and exonic regions, respectively, across replicates, in input (top) and xRIP-seq (bottom) samples in the indicated ELAV pull-down experiments (Flag IP in *elav*$^{FLAG}$ flies, ELAV IP in wild-type flies) and the control experiment (Flag IP in wild-type flies).

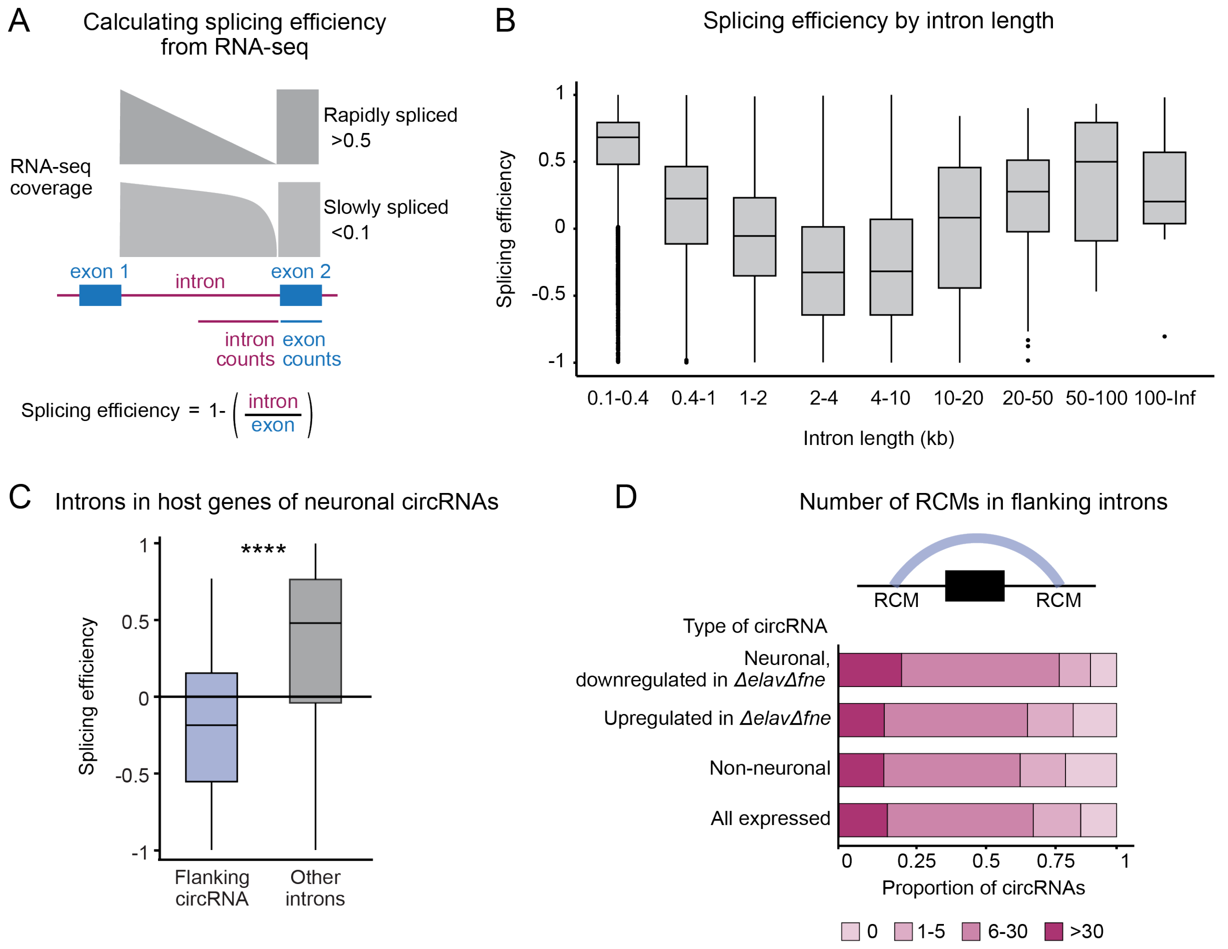


**Figure S4. circRNA biogenesis is associated with poor splicing efficiency. Related to Figure 4.**
(A) Schematic representation of how splicing efficiencies were calculated. Nascent-seq data from (Khodor et al., 2011) were mapped. Reads that aligned to the last 15 nucleotides of the intron (15 nt of the intron 3' end), and reads that aligned to the first 15 nucleotides of the downstream exon (15 nt of the exon 5' end) were quantified. The differential read count in the intron region compared to the exon region was calculated. This value was subtracted from 1 to obtain the splicing efficiency. Values close to 1 represent a high splicing efficiency ("fast splicing").
(B) Box plot showing the distribution of splicing efficiency of introns of different length ranges. The median splicing efficiency for each intron length category is represented by the horizontal line within each box, while the whiskers indicate the range of splicing efficiencies observed. Outliers are shown as individual dots.
(C) Splicing efficiency in genes that produce neuronal circRNAs, comparing introns that flank the neuronal circRNA's BSJ to all other introns of the same gene. ****p<0.0001 (one-tailed Welch's t-test).
(D) For each type of circRNA, proportion of circRNAs whose flanking introns contain reverse complementary match sequences (RCMs). Number ranges indicate RCMs per intron.
